# Ppx1 putative exopolyphosphatase is essential for polyphosphate accumulation in *Lacticaseibacillus paracasei*

**DOI:** 10.1101/2023.12.22.573107

**Authors:** Daniela Corrales, Cristina Alcántara, Manuel Zúñiga, Vicente Monedero

## Abstract

The linear polymer polyphosphate (poly-P) is present across all three domains of life and serves diverse physiological functions. The enzyme polyphosphate kinase (Ppk) is responsible for poly-P synthesis, whereas poly-P degradation is carried out by the enzyme exopolyphosphatase (Ppx). In many *Lactobacillaceae*, the Ppk-encoding gene (*ppk*) is found clustered together with two genes encoding putative exopolyphosphatases (*ppx1* and *ppx2*) each having different domain compositions, with the gene order *ppx1*-*ppk*-*ppx2*. However, the specific function of these *ppx* genes remains unexplored. An in-frame deletion of *ppx1* in *Lacticaseibacillus paracasei* BL23 resulted in bacteria unable to accumulate poly-P, whereas disruption of *ppx2* had no effect on poly-P synthesis. Expression of *ppk* was not altered in the Δ*ppx1* strain, and poly-P synthesis in this strain was only restored by expressing *ppx1* in *trans*. Moreover, no poly-P synthesis was observed when *ppk* was expressed from a plasmid in the Δ*ppx1* strain. Purified Ppx2 exhibited *in vitro* exopolyphosphatase activity, whereas no *in vitro* enzymatic activity could be demonstrated for Ppx1. This observation corresponds with the absence in Ppx1 of conserved motifs essential for catalysis found in characterized exopolyphosphatases. Furthermore, assays with purified Ppk and Ppx1 evidenced that Ppx1 enhanced Ppk activity. These results demonstrate that Ppx1 is essential for poly-P synthesis in *Lc. paracasei* and have unveiled, for the first time, an unexpected role of Ppx1 exopolyphosphatase in poly-P synthesis.

**Importance:** Poly-P is a pivotal molecular player in bacteria, participating in a diverse array of processes ranging from stress resilience to pathogenesis, while also serving as a functional component in probiotic bacteria. The synthesis of poly-P is tightly regulated, but the underlying mechanisms remain incompletely elucidated. Our study sheds light on the distinctive role played by the two exopolyphosphatases (Ppx) found in the *Lactobacillaceae* bacterial group, of relevance in food and health. This particular group is noteworthy for possessing two Ppx enzymes, supposedly involved in poly-P degradation. Remarkably, our investigation uncovers an unprecedented function of Ppx1 in *Lacticaseibacillus paracasei*, where its absence leads to the total cessation of poly-P synthesis, paralleling the impact observed upon eliminating the poly-P forming enzyme, poly-P kinase. Unlike the anticipated role as a conventional exopolyphosphatase, Ppx1 demonstrates an unexpected function. Our results added a layer of complexity to our understanding of poly-P dynamics in bacteria.

## INTRODUCTION

The synthesis of the inorganic phosphate polymer polyphosphate (poly-P) in bacteria has been linked to processes related to bacterial fitness under stressful conditions (1–3). Additionally, poly-P regulates in certain pathogens various infection-associated processes, such as motility, biofilm formation, invasion and intracellular survival (4–8). In probiotic bacteria, the ability to produce poly-P has attracted interest due to its potential effects on maintaining intestinal homeostasis. It has been demonstrated that poly-P derived from probiotics can be internalized by endocytosis, activating the p38 MAPK pathway and inducing cytoprotective factors such as HSP27 (9, 10). This process leads to a reinforcement of the epithelial barrier that can be of interest for the treatment of several intestinal disorders (11–13).

Despite the extensive research carried out for years on poly-P synthesis, the regulation of its production in bacteria still presents some unclear aspects. Poly-P is synthesized by the action of the enzyme poly-P kinase (Ppk; EC 2.7.4.1), which utilizes ATP for extending the linear chain of phosphate, while the enzyme exopolyphosphate (Ppx; EC 3.6.1.11) hydrolyses poly-P (14). However, it remains unresolved how these enzymes, which are sometimes encoded by adjacent genes organized in an operon, coordinate their opposing activities to modulate intracellular poly-P levels. According to the classical regulatory model, Ppx activity is inhibited by the nucleotide guanosine penta/tetraphosphate ((p)ppGpp), an alarmone that controls the stringent response. This mechanism would allow the kinase to prevail over the phosphatase, leading to poly-P accumulation (15, 16). In fact, in certain bacteria such as *E. coli* and *Pseudomonas aeruginosa*, poly-P synthesis is activated by various stresses, including nutrient starvation (15, 17). However, this model, originally established for *E. coli*, has been proven incorrect as poly-P accumulation does not depend on (p)ppGpp synthesis (18). Nonetheless, poly-P synthesis requires the participation of starvation responsive regulators, such as PhoB or NtrC (15), the stringent-response-associated transcription factor DksA and other stress response factors (e.g. the sigma factors RpoS, RpoE and RpoN) (19, 20).

In several lactobacilli species, the presence of a *ppk* gene has been associated with the formation of intracellular poly-P granules, and the number of these granules is enhanced when the bacteria are cultivated in a phosphate-rich medium (21). The *ppk* genes in lactobacilli are found in an operon with two genes encoding putative exopolyphosphatases (*ppx1* and *ppx2*). The inactivation of *ppk* in *Lacticaseibacillus paracasei* and *Lactiplantibacillus plantarum* resulted in bacteria that were unable to form intracellular granules and showed no detectable poly-P levels (22). Similarly, a *ppk*-deficient strain of *Limosilactobacillus reuteri* exhibited poly-P levels less than half of the wild type (23).

This study aimed to further explore the regulation of poly-P accumulation in lactobacilli by investigating the functions of *ppx1* and *ppx2*. Mutants lacking these genes were constructed in *Lc. paracasei*, their impact on poly-P synthesis was evaluated and the catalytic activities of the proteins encoded by *ppk*, *ppx1* and *ppx2* were assayed.

## MATERIALS AND METHODS

### Bacterial strains and culture media

Lactobacilli strains used in this work are listed in Table 1. They were routinely grown in MRS medium (BD-Difco) at 37 °C under static conditions. For poly-P production experiments, strains were grown in MEI medium (5 g/l yeast extract, 5 g/l tryptone, 4 g/l K_2_HPO_4_, 5 g/l KH_2_PO_4_, 0.2 g/l MgSO_4_·7H_2_O, 0.05 g/l MnSO_4_, 1 ml/l Tween 80, 0.5 g/l cysteine and 5 g/l glucose). A similar medium with low phosphate (LP-MEI) was formulated by excluding PO ^-3^ salts.

**Table 1.**
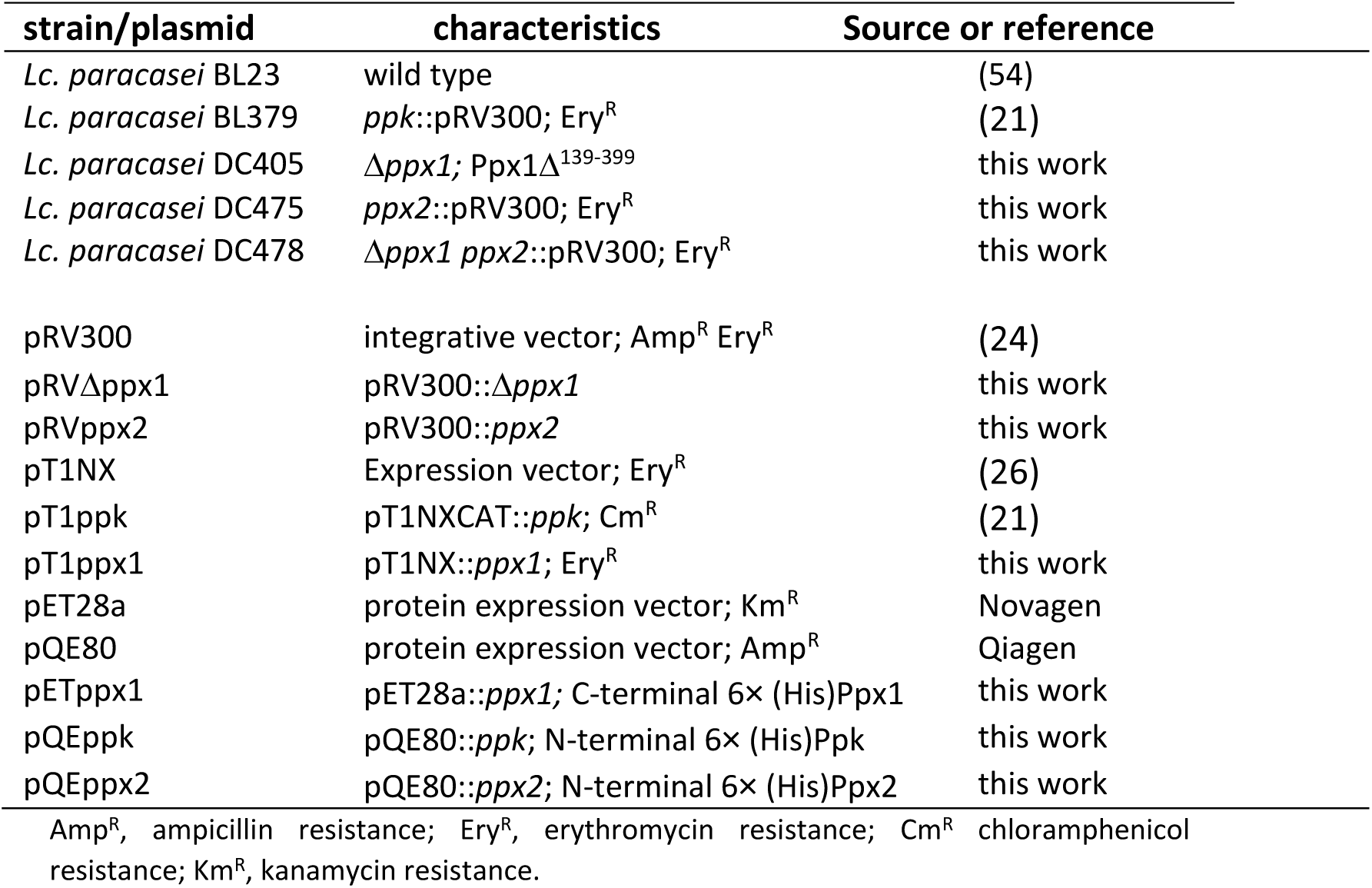
Lactobacilli strains and plasmids used in this work.

*E. coli* NZYStar (*end*A1 *hsd*R17(r_k_-, m_k_+) *sup*E44 *thi-*1 *rec*A1 *gyr*A96 *rel*A1 *lac*[F′ *pro*A^+^B^+^ *lac*I^q^ *ΔlacZ*M15:Tn10(Tc^R^)]) chemically competent cells (NZYtech) were used as an intermediate host for cloning purposes and they were grow in LB medium at 37 °C under agitation. *E. coli* BL21(DE3) (*dcm ompT hsdS*(r_B_-, m_B_-) *gal* λ(DE3)) was used as an expression host for protein purification. *Lactococcus lactis* MG1363 was also used as a host for construction of pT1NX-derived plasmid and it was grown in M17 medium containing 0.5% glucose at 30 °C. When required, erythromycin or chloramphenicol were added at 5 μg/ml for lactic acid bacteria. Ampicillin (100 μg/ml) or kanamycin (25 μg/ml) were used for *E. coli*.

### Construction of strains mutated in *ppx1* and *ppx2*

Two 0.5 kb DNA fragments from *Lc. paracasei ppx1* (LCABL_27830) were obtained by PCR amplification with Phusion High-Fidelity DNA Polymerase (Thermo) and oligonucleotide pairs Ppx1-1/ppx1-2 and ppx1-3/ppx1-4, respectively (Table 2). External primers (ppx1-2 and ppx1-4) had *Eco*RI and *Hin*dIII restriction sites, respectively, at their 5’-ends. Internal primers (ppx1-2 and ppx1-3) overlapped by 20 bp at their ends, thus allowing their fusion by PCR. The resulting 1 kb product carried an in-frame deletion in the *ppx1* reading frame that resulted in the codification of a mutant Ppx1 protein lacking amino acids 139 to 399. The PCR fusion product was digested with *Eco*RI and *Hin*dIII and ligated to pRV300 (24) digested with the same enzymes, and transformed in *E. coli* NZYStar following the procedure recommended by the supplier. The resulting plasmid, pRVΔppx1 (Table 1), was verified by sequencing and used to transform *Lc. paracasei* BL23 strain by electroporation (25) with a Gene Pulser apparatus (Bio-Rad). Integrants at the *ppx1* locus were selected on MRS plates supplemented with erythromycin. One of such integrants was grown for approximately 150 generations in liquid MRS medium without antibiotic pressure and isolates in which a second recombination led to plasmid excision and loss of antibiotic resistance were selected by replica plating. Isolates in which the second recombination led to exchange of *ppx1* by the Δ*ppx1* variant were identified by PCR with appropriate oligonucleotides spanning *ppx1* and confirmed by sequencing. One isolate was selected for subsequent studies (*Lc. paracasei* DC405 strain).

**Table 2.**
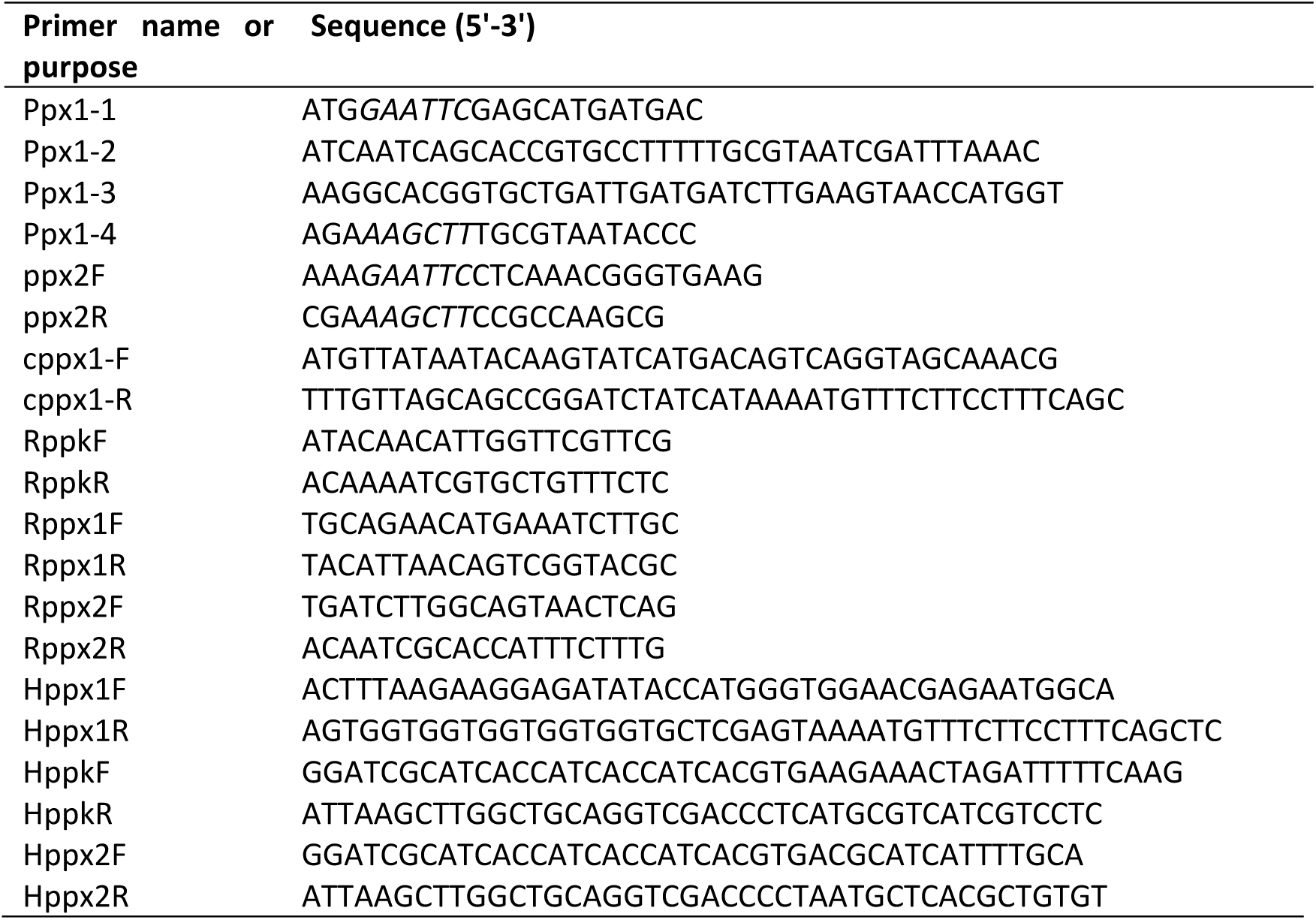
Primers used in this study.

Gene *ppx2* (LCABL_27810) was mutated by insertional inactivation. For this purpose, a 0.4 kb *ppx2* internal fragment was amplified with oligonucleotides ppx2F and ppx2R carrying *Eco*RI and *Hin*dIII restriction sites at their respective 5’-ends (Table 2). The fragment was digested with these enzymes, ligated to pRV300 and transformed into *E. coli* NZYStar, giving pRVppx2. Electroporation of this plasmid into *Lc. paracasei* BL23 resulted in *ppx2* disruption (strain DC475). The pRVppx2 plasmid was also used to transform the DC405 strain, resulting in the double mutant strain DC478 (Δ*ppx1 ppx2*::pRV300).

A plasmid expressing *ppx1* was constructed by amplifying *ppx1* with Phusion High-Fidelity DNA Polymerase and oligonucleotides cppx1-F and cppx1-R (Table 2). The PCR product was cloned into pT1NX (26) digested with BglII and SpeI by Gibson assembly with the GeneArt Gibson Assembly HiFi kit (Invitrogen) following the recommendations of the supplier. The obtained pT1ppx1 plasmid carried *Lc. paracasei ppx1* under the control of the constitutive P1 promoter and was confirmed by sequencing.

### Poly-P isolation and quantification

Poly-P was isolated from bacterial cell pellets (10 ml cultures) by sodium hypochlorite extraction as previously described (21). Poly-P obtained from these bacteria was dissolved in water and quantified by fluorescence with 4′,6′-diamino-2-phenylindole (DAPI; 10 μM in 50 mM Tris-HCl pH 7.5, 50 mM NaCl buffer) (27) in a Clariostar plate reader (Ex. 425 nm, Em. 550 nm). Standard curves of poly-P were built with different amounts of poly-P and they were quantified with DAPI. The standard curves were calibrated by determining total phosphate. To this end, poly-P was subjected to acid hydrolysis (1 vol of 2M HCl, 95 °C, 15 min, followed by neutralization with 1 vol 2M NaOH) and total phosphate was determined with the Biomol Green kit (Enzo Life Sciences). Results are then expressed as nmol phosphate normalized to the amount of bacterial cells expressed as OD at 595 nm.

### Poly-P electrophoresis and staining

The poly-P fractions were analysed in 8% polyacrylamide gels containing 8 M urea and 1X Tris-borate-EDTA (TBE) as running buffer. After electrophoresis, the gels were stained with toluidine blue (28). Poly-P granules in *Lc. paracasei* strains were evidenced by Neisser staining: bacterial cell pellets from liquid cultures were smeared on microscope slides, air dried and stained with a solution of 0.1% methylene blue, 5% acetic acid, 5% ethanol mixed (1:2) with a solution of 0.33% crystal violet in 10% ethanol for 1 min. After rinsing with water, the bacteria were contrasted with 0.3% chrysoidin G in water and observed under microscope at 100X.

### Determination of gene expression by RT-qPCR

Total RNA was isolated from bacteria grown in MEI or LP-MEI with TRIzol (Invitogene) as described previously (29). Quality and concentration of the RNA samples were subsequently evaluated by using the Experion automated electrophoresis system (Bio-Rad). Samples with 23S/16S ratios lower than 0.85 were discarded. RNA was treated with the Ambion Turbo DNA-free™ kit (Applied Biosystems) and cDNA was synthesized from 5 µg RNA with the SuperScript VILO cDNA synthesis kit (Invitrogen). Real-time qPCR was performed using a LightCycler®480 Real-Time PCR system (Roche Diagnostics, USA) with the LightCycler® 480 SYBR Green I Master Mix (2X, Roche) and the primer pairs RppkF/RppkR for *Lc. paracasei* BL23 *ppk* gene (LCABL_27820), Rppx1F/Rppx1R for *ppx1* and Rppx2F/Rppx2R for *ppx2* (Table 2). The housekeeping genes *recA*, *fusA*, *pyrG* and *mutL* were used as reference (30). The cycling conditions were as follows: 95 °C for 5 min, followed by 40 cycles of three steps consisting of denaturation at 95 °C for 10 s, primer annealing at 55 °C for 10 s, and primer extension at 72 °C for 10 s. Relative gene expression was determined with the tools implemented in the Relative Expression Software Tool (31).

### Expression and purification of His-tagged proteins

The coding regions for *Lc. paracasei* BL23 *ppx1* (LCABL_27830), *ppk* (LCABL_27820) and *ppx2* (LCABL_27810) genes were amplified by PCR with the oligonucleotide pairs Hppx1F/Hppx1R, HppkF/HppkR and Hppx2F/Hppx2R (Table 2), respectively, using the Phusion High-Fidelity DNA Polymerase (Thermo Scientific). The purified PCR fragments were cloned into expression plasmids pET28a, digested with *Nco*I/*Xho*I (for *ppx1*) or pQE80, digested with *Sma*I-*Bam*HI (for *ppk* and *ppx2*), with the GeneArt Gibson Assembly HiFi kit (Invitrogen), in order to express proteins with C-terminal or N-terminal 6X(His) tags, respectively. Ligation mixtures were used to transform *E. coli* NZYStar and, after sequencing, the constructed plasmids were transferred to *E. coli* BL21(DE3). For protein expression, cultures (0.5 l) were grown in LB broth at 37°C with shaking (200 r.p.m.) to an OD_595_ 0.5. Protein expression was induced by adding 0.3 mM IPTG and incubation at 27°C for 3 h (Ppx1), 0.3 mM IPTG and overnight incubation at 16°C (Ppk) or 0.3 mM IPTG and 3h incubation at 27°C (Ppx2). After centrifugation, bacterial pellets were washed with 200 ml of 20 A buffer (Tris-HCl 50 mM [pH 7.4], NaCl 500 mM), and stored at -80°C until use. Cells were thawed on ice and resuspended in 30 ml of A buffer supplemented with 1 mM phenylmethylsulfonyl fluoride. The cell suspension was sonicated at 200 W for four cycles of 60 s followed by 60 s of cooling with a Labsonic V sonicator (B. Braun). The cell debris was removed by centrifugation at 12,000 × *g* for 15 min at 4°C and the supernatant filtered through a 0.45 μm pore size Filtropur S membrane (Sarstedt). The cleared extract was directly loaded onto a HisTrap FF crude column (GE Healthcare Life Sciences) equilibrated with buffer A and coupled to an Äktaprime FPLC system (Cytiva). After the passage of the sample, the column was washed with 10 ml of A buffer and the proteins were eluted with an imidazole gradient (40 mM to 2 M) in A buffer. Fractions were analysed by SDS-PAGE and those containing purified protein were pooled and subjected to buffer exchange (50 mM MES pH 6.5, 100 mM KCl, 1 mM MgCl_2_ for Ppx1 and Ppx2, whereas KCl was increased to 200 mM for Ppk) using an Amicon Ultra 10K centrifugation device (Merck Millipore). Glycerol was added to 20% (v/v) final concentration and the recombinant proteins were kept at -80°C.

### Enzymatic activities

Exopolyphosphatase activity was measured in 150 μl of 50 mM MES pH 5.5, 100 mM KCl, 1 mM MgCl_2_, poly-P isolated from *Lc. paracasei* BL23 (equivalent to 2 nmol of Pi) and different amounts (0.3 to 0.7 μg) of purified 6x(His)Ppx1 or 6x(His)Ppx2. The mixture was incubated at 37°C and 10 μl aliquots were withdrawn at different time intervals and mixed with 140 μl of Biomol Green reagent (Enzo Life Sciences) to determine phosphate release. Poly-P kinase activity was determined in 100 μl of 50 mM HEPES-KOH pH 7, 5 mM MgCl_2_, 40 mM ammonium sulphate, 2 mM ATP, 0.05 nmol of poly-P (as inorganic phosphate), 0.2 μg of 6x(His)Ppk, 2.5 mM phospho*enol*pyruvate (PEP) and 2U of pyruvate kinase (Sigma). PEP and pyruvate kinase in the reactions were used to regenerate ATP and to avoid the accumulation of ADP, which may have inhibitory activity on Ppk (32). When required, 0.2 μg of *Lc. paracasei* 6x(His)Ppx1 or BSA were added to the reactions. The lack of poly-P kinase activity in 6x(His)Ppx1 or BSA was also tested in reaction in which these proteins were added in the absence of 6x(His)Ppk. The mixtures were incubated at 37°C and at different time intervals, 10 μl aliquots were mixed with 10 μl of 500 mM EDTA pH 8 to stop the reaction. Samples were kept on ice until poly-P formation was quantified by DAPI fluorescence in a Clariostar fluorescence plate reader.

## RESULTS

### Poly-P synthesis and degradation is linked to phosphate availability and growth phase in *Lacticaseibacillus paracasei* BL23

It has been previously shown that growth in phosphate-rich medium favours the formation of poly-P intracellular inclusions in many lactobacilli, including *Lc. paracasei* (21). Here we show that poly-P amounts were around five-fold higher in *Lc. paracasei* BL23 cells grown in phosphate-rich compared to low phosphate medium at late exponential growth (6h of culture, OD ∼2.5; Figure 1A). In stationary phase cells grown in MEI (high phosphate; 16h of culture, OD ∼3.1), poly-P contents decreased to that of cells grown in LP-MEI at the late exponential phase. This difference in poly-P levels between exponential and stationary cells was also observed when bacteria were cultivated in LP-MEI (Figure 1A). When the isolated poly-P was analysed by acrylamide-urea electrophoresis, differences in chain lengths were also evidenced in cells grown in MEI. The poly-P fraction obtained from exponentially growing bacteria displayed a higher content of long chain poly-P than that obtained from stationary phase cells (Figure 1B). This difference in poly-P length was not observed in poly-P isolated at different growth stages from cells grown in LP-MEI (Figure 1B). To test whether differences in poly-P contents could be related to differential expression of the *ppx-ppk* gene cluster, expression of *ppx1*, *ppk* and *ppx2* was determined (Figure 2). We showed that expression of *ppx* and *ppk* genes was slightly reduced in low phosphate grown cells compared to those grown in high phosphate conditions both in late exponential and stationary phase. This result suggests that phosphate availability has a moderate effect on *ppx*/*ppk* gene expression.

**Figure 1.**
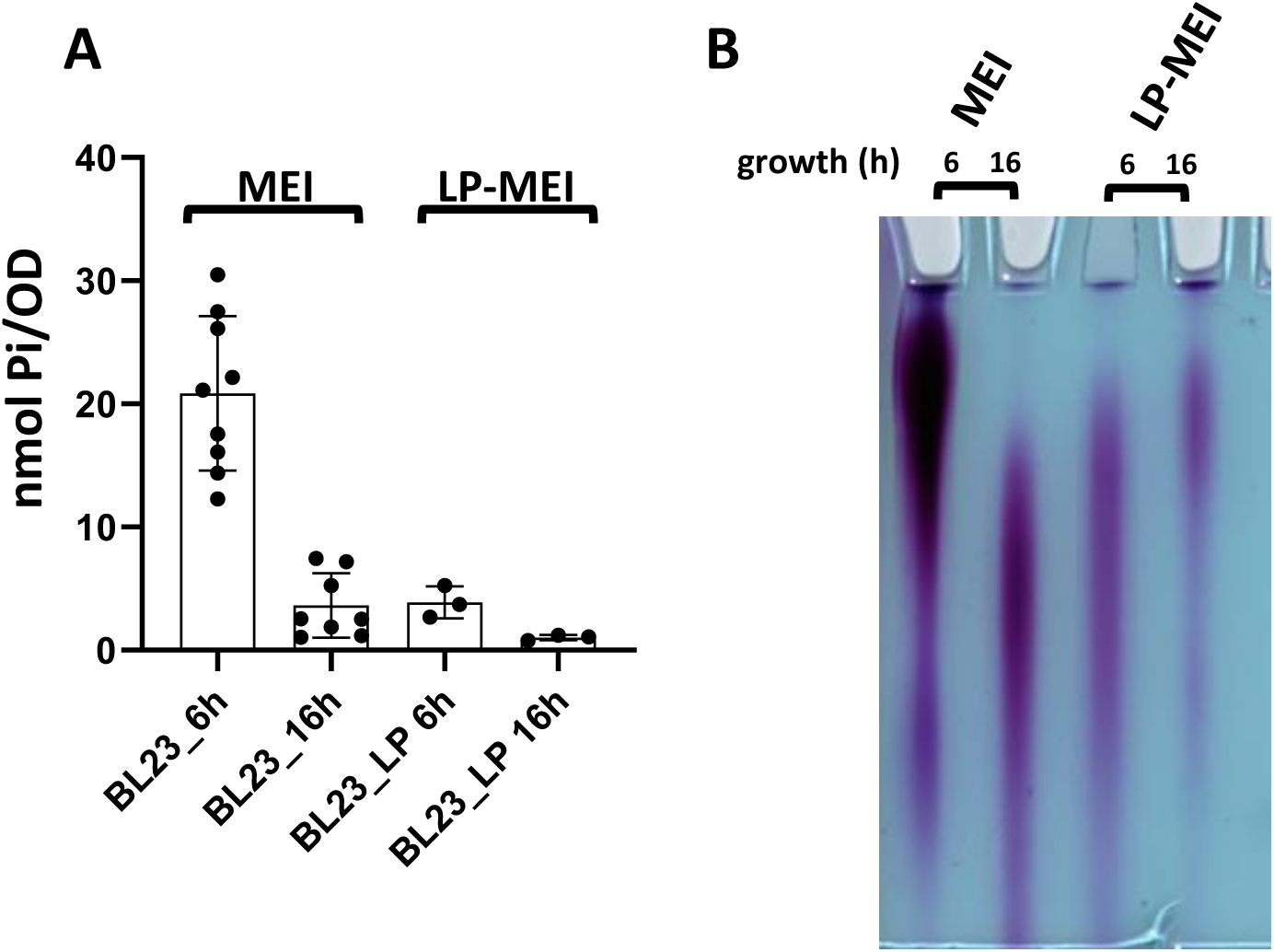
Poly-P synthesis in *Lc. paracasei*. (A) levels of poly-P as nmol Pi/OD in wild-type *Lc. paracasei* BL23 at different times of growth in MEI medium (high phosphate 6h; OD∼2.5 and 16h; OD∼3.1) and LP-MEI (low phosphate; LP. 6h; OD∼1 and 16h; OD∼1.6); (B) analysis of poly-P by acrylamide electrophoresis in *Lc. paracasei* BL23 grown under the same conditions as in (A). Each lane contains the poly-P extracted from bacterial cells equivalent to 4 OD units.

**Figure 2.**
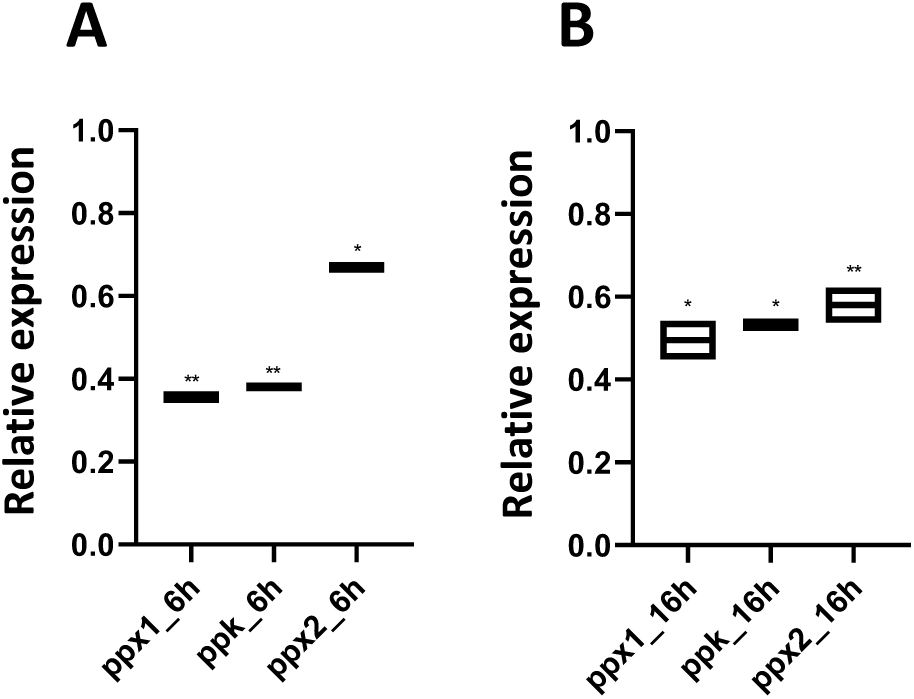
Expression of *ppk* and *ppx* genes. Relative expression (fold-change) of *ppk*, *ppx1* and *ppx2* in *Lc. paracasei* BL23 cells grown in LP-MEI compared to cells grown in MEI. (A) RNA isolated from cells grown for 6 h. (B) RNA isolated from cells grown for 16 h. Asterisks indicate significant differences respect to reference conditions, * p<0.05, ** p< 0.005.

### A deletion in the *ppx1* gene encoding a putative exopolyphosphatase results in the absence of poly-P accumulation

The *ppk* gene from many lactobacilli, including *Lc. paracasei*, is clustered with *ppx1* and *ppx2*, presenting an operon structure *ppx1*-*ppk*-*ppx2* ((21); Figure 3A). Stop and start codons for *ppx1*/*ppk* and *ppk*/*ppx2* gene pairs are overlapped, respectively, indicating a strong translational coupling in the synthesis of the three enzymes. Ppx1 has a protein architecture similar to other exopolyphosphatases belonging to the GppA/Ppx family, such as that of *E. coli*, with an N-terminal catalytic domain belonging to the acetate and sugar kinase/Hsp70/actin (ASKHA) superfamily and a C-terminal HDc domain present in metal-dependent phosphohydrolases (33). The exopolyphosphatase Ppx2 shares low amino acid identity (15%) with Ppx1. Ppx2 contains the ASKHA domain whereas it lacks the HDc domain.

**Figure 3.**
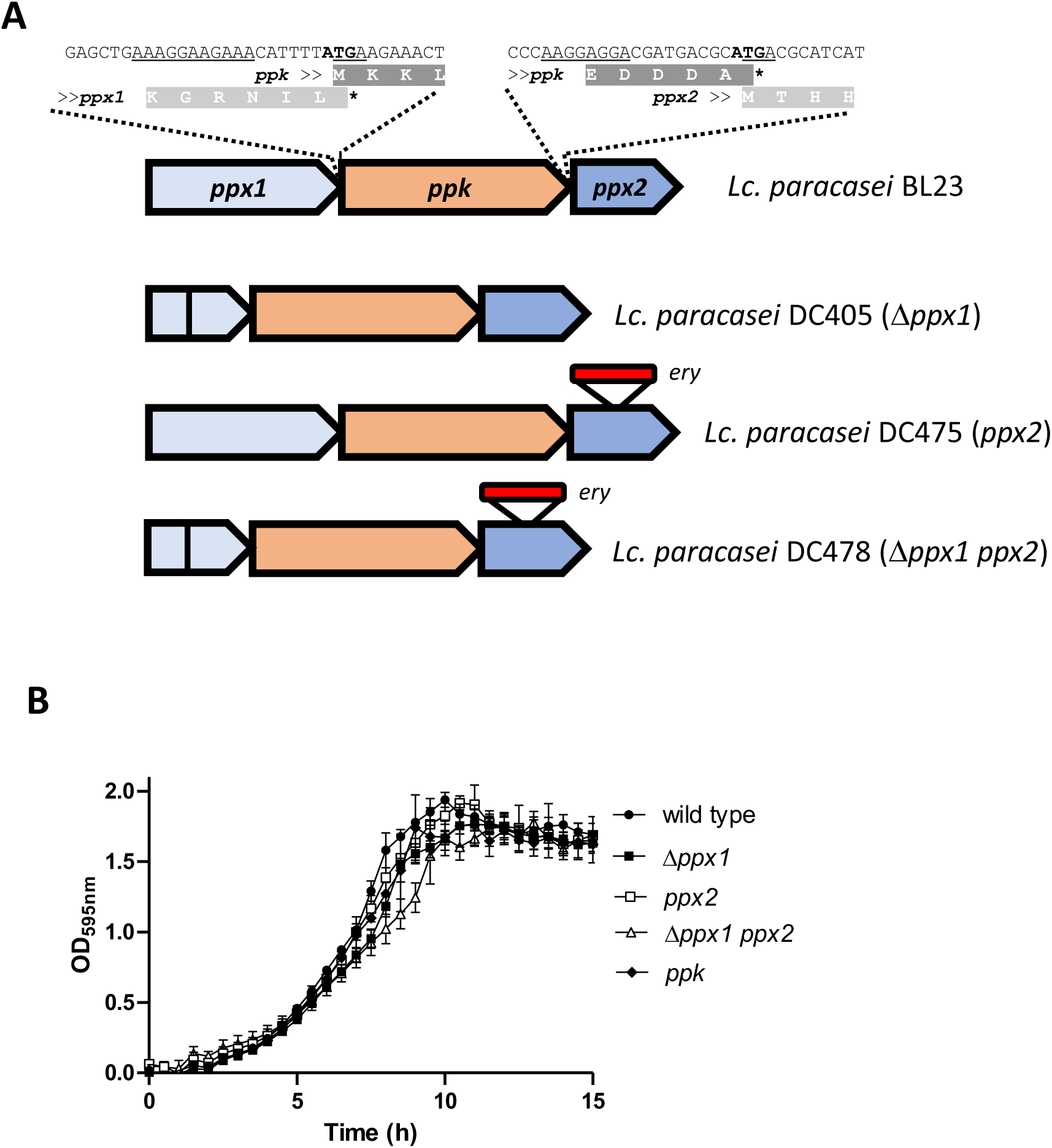
Mutated strains in the *ppx1*-*ppk*-*ppx2* operon. (A) Schematic representation of the *ppx1*-*ppk*-*ppx2* operon in *Lc. paracasei* BL23. The overlapping start and stop codons for *ppx1*-*ppk* and *ppk*-*ppx2* are shown. Underlined sequences are Shine-Dalgarno sites. The different operon structure in the constructed *ppx* mutants is also shown for DC405 (in-frame deletion in *ppx1*), DC475 (plasmid integration in *ppx2*) and DC478 (plasmid integration in *ppx2* in the Δ*ppx1* mutant) strains. (B) Growth curves of different *Lc. paracasei* BL23 derivatives. The strains were incubated in MEI medium at 37 °C and growth was monitored as OD at 595nm.

In order to gain insight about the function of these Ppx enzymes during poly-P accumulation, an *Lc. paracasei* strain with an in-frame deletion in *ppx1* was constructed which resulted in deletion of amino acids 139 to 399 in Ppx1 and therefore lacked the putative enzyme active site and a substantial part of its amino acid sequence. A strain with a disruption of *ppx2* was also obtained and a double mutant strain was created by interrupting *ppx2* in the Δ*ppx1* strain (Figure 3A). Similar to a previously described *ppk*-deficient strain from *Lc. paracasei* BL23 (21), the different introduced mutations did not show a strong impact on growth under the tested conditions (Figure 3B). Surprisingly, no poly-P could be detected in the Δ*ppx1* strain (Figure 4A). However, the poly-P content and size distribution in the *ppx2* strain were similar to those found in the wild-type strain (Figure 4B). In addition, strain *ppx2* followed the same pattern of poly-P accumulation than the wild-type strain, being dependent on the growth phase. Inspection of intracellular granules confirmed poly-P measurements (Figure 4C): poly-P granules were detected in wild-type and *ppx2* strains whereas they were absent in the *ppk* mutant, the *ppx1* deleted strain and the double mutant Δ*ppx1 ppx2*.

**Figure 4.**
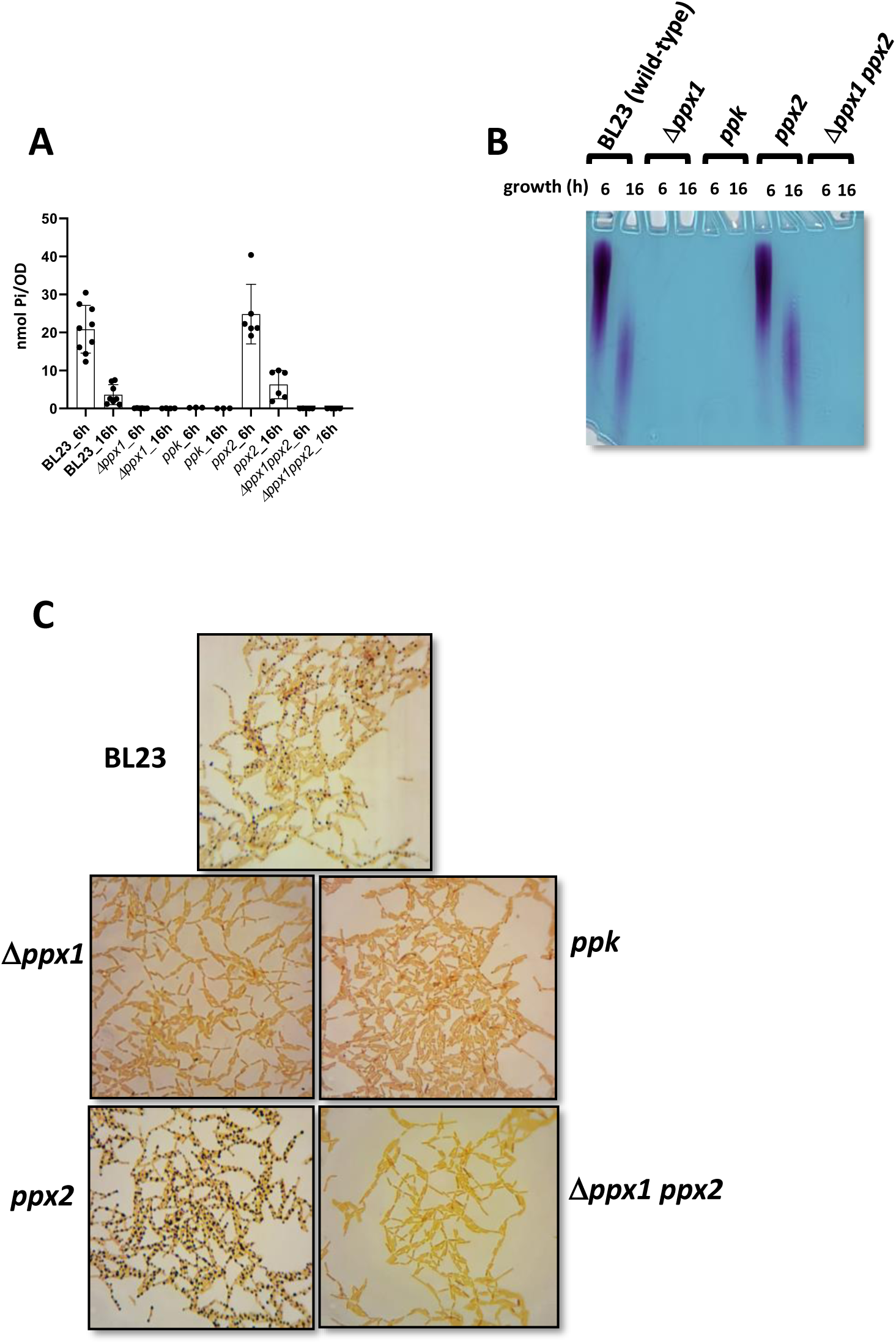
Effect of *ppx* mutations on poly-P synthesis. (A) levels of poly-P as nmol Pi/OD in *Lc. paracasei* BL23 (wild-type) and mutant strains in *ppx1*, *ppx2* or *ppk* at different times of growth in MEI (high phosphate 6h; OD∼2.5 and 16h; OD∼3.1) and LP-MEI (low phosphate 6h; OD∼1 and 16h; OD∼1.6) media; (B) analysis of poly-P by acrylamide electrophoresis in each strain grown under the same conditions as in (A). Each lane contains the poly-P extracted from bacterial cells equivalent to 4 OD units. The wild-type (BL23) data are the same measurements shown in Figure 1 (n=9), derived from independent experiments performed at different days. For the rest of strains n=3-6. (C) Poly-P granule formation in *Lc. paracasei*. Bacterial cells from different strains were grown in MEI for 16h, stained by Neisser staining and visualized at 100X.

### Ppx1 is essential for poly-P synthesis in *Lc. paracasei*

In order to investigate the lack of poly-P synthesis after *ppx1* deletion we measured the levels of *ppk* gene expression in the Δ*ppx1* strain, showing that there were no relevant differences compared to the wild-type strain (Figure 5A). Despite of similar *ppk* expression between strains, the lack of poly-P synthesis as a consequence of the *ppx1* deletion could still rely on differences in Ppk expression caused by the upstream changes in *ppx1*. Refuting this hypothesis, transformation of the Δ*ppx1* strain with a plasmid carrying wild-type *ppx1* restored poly-P synthesis and granule formation (Figure 5B, C and D). On the contrary, after transformation with a plasmid expressing *ppk*, which in the *ppk* mutant was able to reestablish poly-P production and granule formation, no poly-P was detected (Figure 5B and C). According to this, expression of plasmid-carried *ppk* in the Δ*ppx1* mutant resulted in no poly-P granule formation (Figure 5D). Although complementation of Δ*ppx1* and *ppk* mutations by their respective wild-type alleles was observed, in the complemented strains the levels and pattern (chain length distribution) of poly-P never reached those of the wild-type. It has to be considered that in these experiments *ppk* and *ppx1* genes were fused to a constitutive promoter and expressed from multicopy plasmid, for which they were probably not subject to the same regulation as the chromosomal genes.

**Figure 5.**
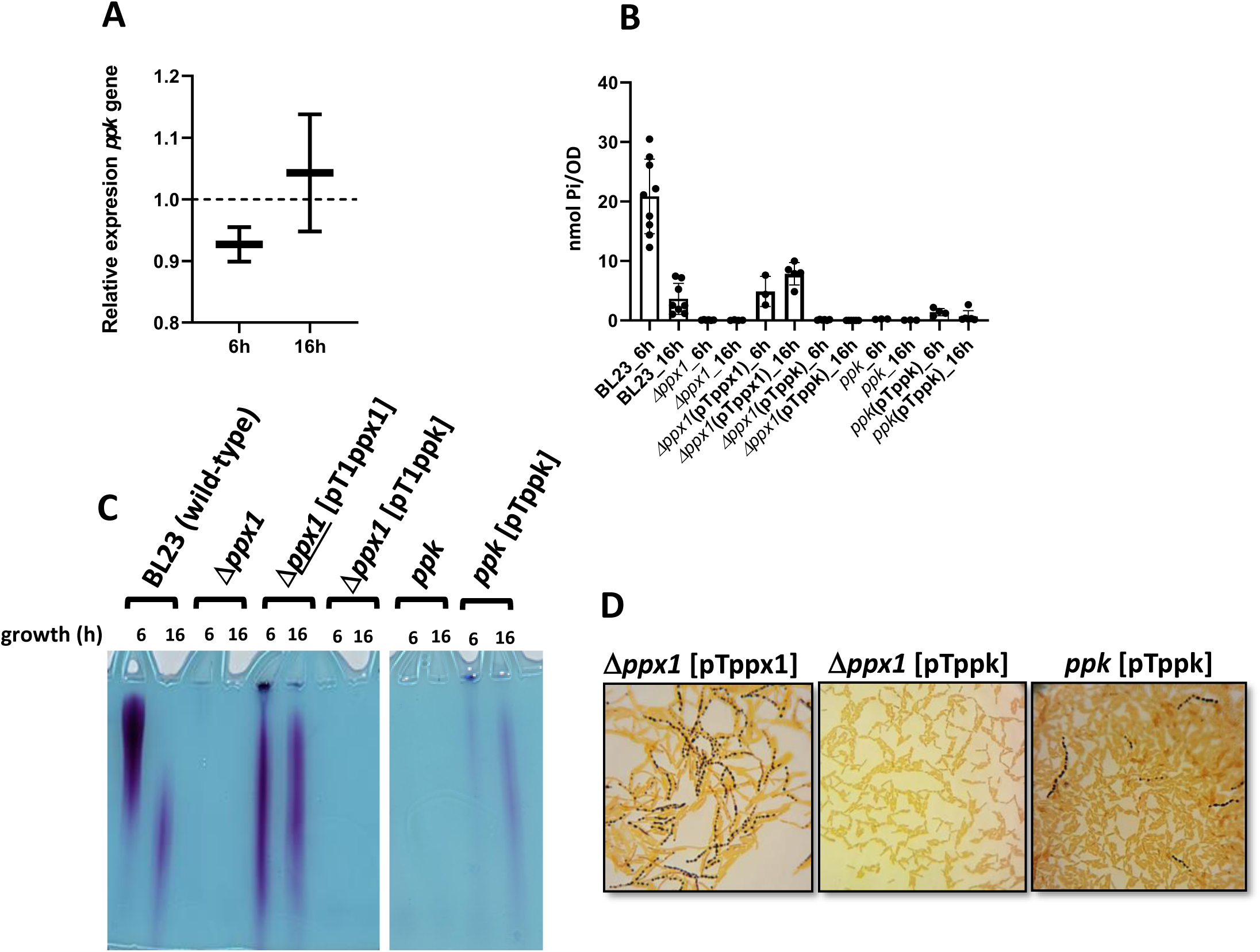
Contribution of Ppx1 to Poly-P biosynthesis in *Lc. paracasei.* (A) Relative expression of *ppk* in a 1*ppx1* strain. Expression of *ppk* was quantified by qPCR at different culture times in MEI in the *Lc. paracasei* 1*ppx1* strain. Data are fold-changes relative to wild-type cells. (B) Levels of poly-P as nmol Pi/OD in *Lc. paracasei* BL23 (wild-type) and mutant strains in *ppx1*, or *ppk* and their respective variants carrying plasmids bearing *ppx1* or *ppk* at different times of growth in MEI (high phosphate 6h; OD∼2.5 and 16h; OD∼3.1) and LP-MEI (low phosphate 6h; OD∼1 and 16h; OD∼1.6) media; (C) analysis of poly-P by acrylamide electrophoresis in each strain grown under the same conditions as in (B). Each lane contains the poly-P extracted from bacterial cells equivalent to 4 OD units. The wild-type (BL23) data are the same measurements shown in Figure 1 (n=9), derived from independent experiments performed at different days. For the rest of strains n=3-6. (D) Poly-P granules staining. Bacterial cells from different *Lc. paracasei* strains were grown in MEI for 16h, stained by Neisser staining and visualized at 100X.

### Exopolyphosphatase and poly-P kinase activities of *Lc. paracasei* BL23 enzymes

As already mentioned, the *Lc. paracasei* Ppx1 protein belongs to the *E. coli* GppA/Ppx prototype. However, a close inspection of the sequence of Ppx1 from *Lactobacillaceae* revealed mutations in key amino acids that have been identified in Ppx proteins as crucial for their catalytic activities (Figure 6). Ppx proteins from the GppA/Ppx family carry glycine-rich phosphate-binding loops (P-loop motifs) in their catalytic N-terminal domains that are important for accommodating phosphate moieties from the poly-P polymer (33). The first P-loop, with a consensus sequence DXGS[N/Y]S, was absent in the *Lactobacillaceae* Ppx1 proteins, whereas the second P-loop, which displays a consensus sequence [D/E]XGG[G/A]SXE, largely deviated from it (DISSGSVE for *Lc. paracasei* Ppx1, showing a replacement of the highly conserved central glycine residues by serine; Figure 6). For some lactobacilli species, sequence conservation at this level was very poor, showing a complete lack of conservation for the aspartic and glutamic residues present at the boundaries of the P-loop 2, as was the case for Ppx1 from *Lp plantarum* (WP_103851489.1), *Levilactobacillus brevis* (AYM03703.1), *Limosilactobacillus reuteri* (MCH5356744.1) or *Lactobacillus acidophillus* (MCT3601606.1). The catalytically important general acid-base glutamic residue found in GppA/Ppx, situated proximal to P-loop 2 (E121 in *E. coli* Ppx), was identified in certain Ppx1 proteins within the *Lactobacillaceae* family. However, it was notably absent in others (e.g. *Lm. reuteri* or *L. acidophillus*). This residue is aligned through an arginine (R93 in *E. coli* Ppx) to facilitate the activation of the nucleophile water, promoting its attack on the terminal phosphate in poly-P (34, 35). Nevertheless, this crucial arginine residue was not observed in Ppx1 from *Lactobacillaceae*. On the contrary, the Ppx2 proteins from *Lc. paracasei*, but also from other *Lactobacillaceae* family members carrying *ppx1*-*ppk*-*ppx2* clusters, displayed full conservation of P-loops 1 and 2 (e.g.: DLGSNS and DTGGASTE amino acid sequences, respectively, for *Lc. paracasei* Ppx2) and those key catalytic amino acids (Figure 6).

**Figure 6.**
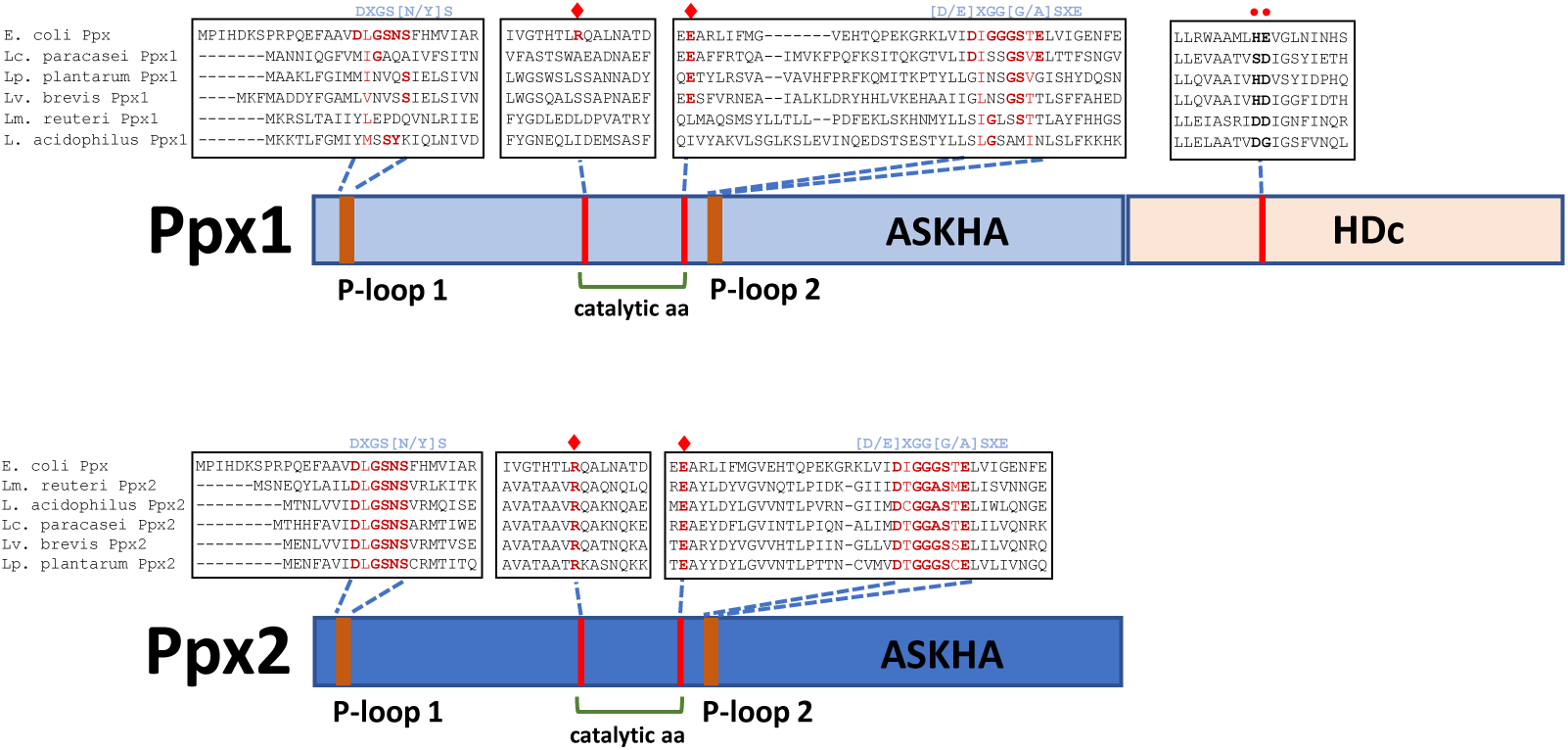
Schematic representation of Ppx1 and Ppx2. The presence of different protein domains (ASKHA and HDc) is depicted, together with relevant portions of multiple sequence alignments of Ppx1 from *Lc. paracasei* (WP_003606084.1), *Lv. brevis* (AYM03703.1), *Lp. plantarum* (WP_103851489.1), *Lm. reuteri* (MCH5356744.1) and *L. acidophilus* (MCT3601606.1) and from Ppx2 from *Lc. paracasei* (CAQ67832.1), *Lv. brevis* (WP_135367400.1), *Lp. plantarum* (WP_003641094.1), *Lm. reuteri* (KRK50994.1) and *L. acidophilus* (AZN76677.1), compared with Ppx from *E. coli* (WP_001121363.1). The consensus sequences of the conserved P-loop 1 and P-loop 2 in the GppA/Ppx family are shown in blue above the alignments. The catalytic amino acids arginine and glutamic acid are marked with red diamonds. The amino acids characteristic of the HDc domain in C-terminal Ppx1 are marked with red circles. For full protein alignments see Supplementary Figure S2.

To confirm the enzymatic activities of the proteins encoded within the *Lc. paracasei* BL23 *ppx1*-*ppk*-*ppx2* cluster, the three proteins were expressed and purified from *E. coli* as His-tagged proteins (Supplementary Figure S1). In agreement with the presence of all sequence characteristics linked to exopolyphosphatases, recombinant Ppx2 exhibited activity on poly-P from *Lc. paracasei*, releasing 2.54 ± 0.41 nmol of phosphate/min/μg protein. However, no activity could be evidenced for the purified Ppx1 protein (<0.003 nmol of phosphate/min/μg protein; below detection level). Exopolyphosphatase activity from Ppx1 could also not be detected (below detection level) when Ppk was added to the reaction mixture. In contrast to the polyphosphatase activity of Ppx1, poly-P kinase activity was evidenced for the *Lc. paracasei* Ppk enzyme. The poly-P kinase activity of Ppk was enhanced in reactions where the Ppx1 protein was included, which, when tested alone, did not display this activity (Figure 7).

**Figure 7.**
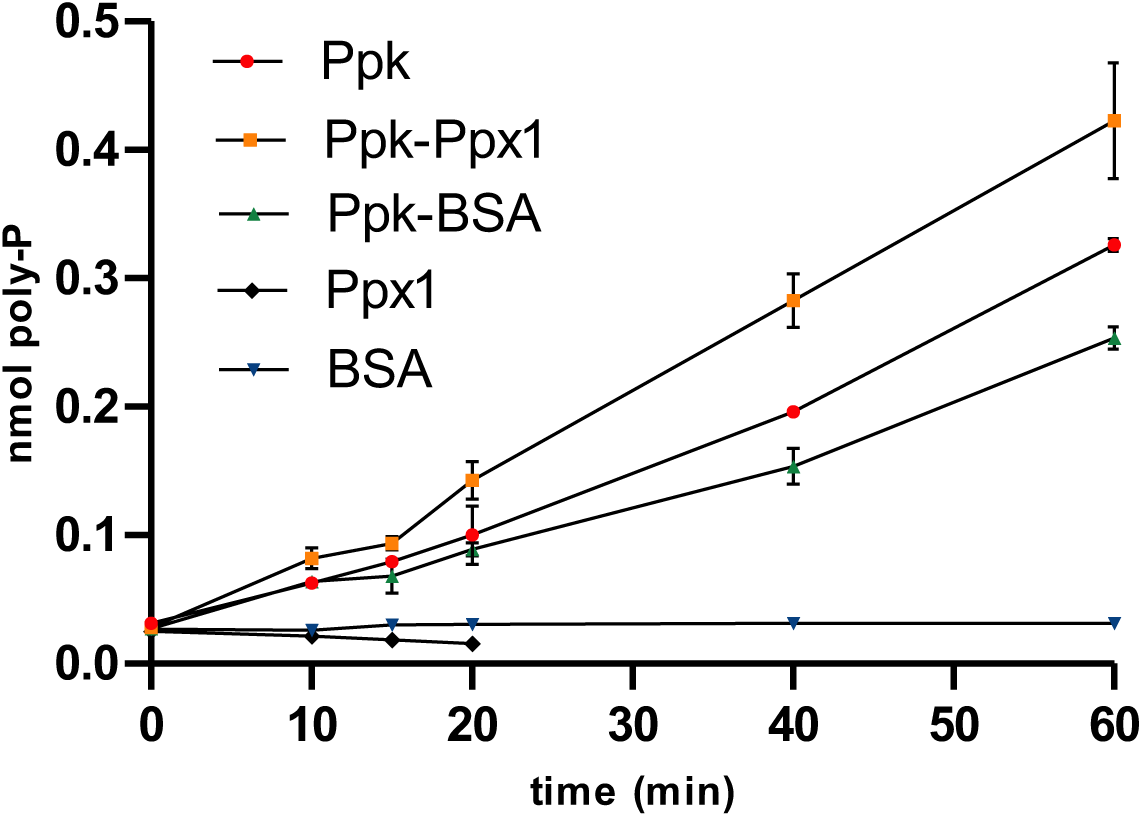
Poly-P kinase activity of *Lc. paracasei* Ppk. Purified 6x(His) Ppk from *Lc. paracasei* BL23 was assayed for poly-P kinase in the absence or presence of 6X(His)Ppx1. BSA was employed as a control unrelated protein and the poly-P kinase activity of the added 6X(His)Ppx1 was also checked.

## DISCUSSION

It has been commonly accepted that poly-P levels in bacteria are controlled by the antagonistic activities of Ppk (synthesis) and Ppx (hydrolysis). However, the model that pointed to (p)ppGpp as a key regulatory molecule inhibiting Ppx activity, and hence triggering poly-P accumulation, has been recently questioned (18). How poly-P synthesis is regulated in lactobacilli is not known. We showed that its levels in *Lc. paracasei* BL23 vary depending on the amount of phosphate in the medium ((21) and results presented here), a fact that has also been reported in *Lacticaseibacillus rhamnosus* CRL 1505 (36). In addition, as described in this work, other researchers reported that an initial accumulation at early growth phases was followed by a decrease in poly-P concentration in *Lc. paracasei* JCM1163 and phosphate-starved cells of this strain also accumulated poly-P after phosphate supply (13). This paralleled poly-P accumulation in *E. coli, P. aeruginosa* or *Acinetobacter johnsonii* after nutritional downshift, which is also transitory (15). The differences in the poly-P chain lengths that were observed here at different growth stages probably reflect the degradation process that occurs along growth after poly-P accumulation at initial stages.

Whether transcription regulation of *ppk* would play a role in controlling poly-P synthesis is controversial. The *Lc. paracasei ppk* and *ppx* messenger levels were higher under high-phosphate conditions, although the three genes (*ppk*, *ppx1* and *ppx2*), which theoretically encode enzymes with opposite functions, followed the same expression pattern. This agrees with the genetic organization in an operon of the gene cluster (Figure 3A). In *Pseudomonas aeruginosa*, *ppk* and *ppx* (which are not located in the same operon) are differentially regulated (37). In this bacterium these two genes do not respond to phosphate availability (do not belong to the *pho* regulon), but their transcription is up- and down-regulated, respectively, upon *phoU* mutation (37), which enhances phosphate uptake and contributes to poly-P accumulation (38). The mutation of *phoU* also increases poly-P synthesis in *E. coli* (39). In contrast*, ppk* expression in *Acinetobacter* (40) or *Streptomyces lividans* (28) was induced upon phosphate starvation. In addition, other studies showed that in *P. aeruginosa*, transcription of *ppx* is controlled by the response regulator PhoB, which binds to its promoter under phosphate limiting conditions (41) and in *Pseudomonas fluorescens ppk* forms part of the *pho* regulon and it is upregulated in response to low phosphate conditions (42). Therefore, transcriptional regulation of *ppk* and *ppx* genes possibly responds to diverse signalling pathways in different bacteria. Whether the *Lc. paracasei ppk* and *ppx* genes are members of the *pho* regulon (i.e. controlled by PhoB) has to be investigated.

If Ppx was a factor limiting poly-P accumulation in *Lc. paracasei*, in the absence of it or when Ppx activity was diminished, an increase in the cell poly-P content would be expected. Therefore, the fact that inactivation of *ppx1* resulted in the complete absence of poly-P in *Lc. paracasei* was surprising. As far as we know, this is the first time that this effect is reported. In *E. coli*, *ppx* deleted strains accumulated slightly more poly-P compared to wild-type cells, suggesting that activation of Ppk is the main mechanism at play for regulating the amount of poly-P and Ppx would only control its levels once Ppk is activated (43). Also, simultaneous deletion of *ppx1* and *ppx2* genes in *Campylobacter jejuni* (both encoding Ppx harbouring ASKHA and HDc domains), only resulted in less than 1.4-fold increase in poly-P (5) and deleting the *Corynebacterium glutamicum ppx2* gene, impacted poly-P concentration by two-fold (44). Finally, deletion of *ppx* in *Pseudomonas putida*, which is located downstream *ppk* but transcribed convergently, reduces poly-P amounts in both exponential and stationary phase cells by 30-40 % (45). Ppx enzymes from *Lactobacillaceae* have not been studied so far and their functions in poly-P metabolism are not known. While Ppx1 resembles typical exopolyphosphatases from many microorganisms, the shorter Ppx2 lacks the C-terminal HDc domain (46) that in *E. coli* Ppx plays a role in the processivity of the enzyme (47). However, *ppx2* disruption in *Lc. paracasei* did not have an effect on poly-P under the tested conditions. Some species of *Actinomycetales* carry two *ppx2* genes in their chromosomes (44, 48), not clustered with *ppk*, and lack *ppx1* homologues and in the green-sulphur bacterium *Chlorobium tepidum*, two different Ppx have been characterized with protein architectures similar to *Lc. paracasei* (1). For some of these characterized enzymes, biochemical data revealed different substrate preferences (e.g. towards (p)ppGpp or poly-P with different chain lengths (1, 44, 48). However, the amino acid differences at critical positions for Ppx activity in *Lc. paracasei* Ppx1 and orthologous proteins of other *Lactobacillaceae* are intriguing. Mutations in specific amino acids at these sites result in complete loss of enzymatic activity in prototype Ppx1-type enzymes from *E. coli* (34), *Helicobacter pylori* (33) or *P. aeruginosa* (49). Thus, our inability to detect any exopolyphosphatase activity with purified Ppx1 under our laboratory conditions raises the question of whether this protein genuinely functions as an exophosphatase acting on poly-P in *Lc. paracasei*.

The molecular mechanisms by which poly-P synthesis is activated are still elusive. The fact that in *Lc. paracasei* Ppx1 was needed to make poly-P was totally unexpected and would reflect a coordinated action of Ppx1 and Ppk by which the Ppk requires Ppx1 for PolyP synthesis *in vivo*. The existence of factors activating the Ppk kinase function has been hypothesized for *E. coli*. In this bacterium, specific amino acid substitutions in Ppk have been isolated that resulted in a marked increase in poly-P accumulation without affecting enzyme activity (43). Strains carrying these *ppk* mutant alleles constitutively synthesized poly-P, although its level was still raised upon nutritional downshift. It was therefore hypothesized that the amino acid changes in these mutant Ppks render enzymes affected in the *in vivo* interaction of Ppk with factors activating the kinase function (43). One working hypothesis for explaining the phenotype of *ppx1* deletion in *Lc. paracasei* would be that, *in vivo*, *Lc. paracasei* Ppk activation requires the direct interaction with Ppx1 or that Ppx1 participates in the molecular signalling that leads to Ppk activation. The observation that the inclusion of Ppx1 in *in vitro* Ppk reactions enhances poly-P synthesis suggests one of these potential scenarios. However, the mechanism by which this likely process could operate *in vivo* and the signal(s) to which it would respond remain subjects for future investigation. Little is known about the ultrastructure of the granules where poly-P is accumulated and where the machinery for making poly-P should be associated. Distinct proteome and biochemical analysis have identified several proteins bound to the poly-P granules, such as the histone-like protein AlgP from *P. aeruginosa* (50) or proteins carrying the positively-charged <u>C</u>onserved <u>H</u>istidine <u>α</u>-helical <u>D</u>omain (CHAD) in *Ralstonia eutropha*, *Magnetospirillum gryphiswaldense* and *P. putida* (51). Diverse Ppk enzymes are also associated with poly-P granules in *R. eutropha* (52), *Caulobacter crescentus* (53) or *P. aeruginosa* (50), but Ppx has not been yet identified in these kind of analyses (50).

Our results showed that, as occurred in other bacteria, the synthesis of poly-P in lactobacilli is tightly regulated. They also revealed an unexpected phenotype for the lack of Ppx1 in *Lc. paracasei*, which is radically different to that described for *E. coli ppx* mutants or for mutants in other species lacking *ppx1* and/or *ppx2*. Regulation of poly-P production may have implications in the functionality of many probiotic strains, where poly-P can be considered a functional factor with consequences on intestinal homeostasis. Furthermore, the question of which are the mechanisms controlling poly-P synthesis remains open and it should be investigated if strains with enhanced capacity to produce this probiotic factor want to be obtained.

## Acknowledgements

1. D. Corrales is grateful for her doctoral fellowship from the Ministerio de Ciencia, Tecnología e Innovación of Colombia (Convocatoria 906 de 2021, Doctorados en el Exterior). This research was partly funded by project AGL2013-40657-R and supported by the Severo Ochoa Accreditation CEX2021-001189-S from the Ministry of Science, Innovation and Universities from Spain.

---

**Supplementary Figure S1.**
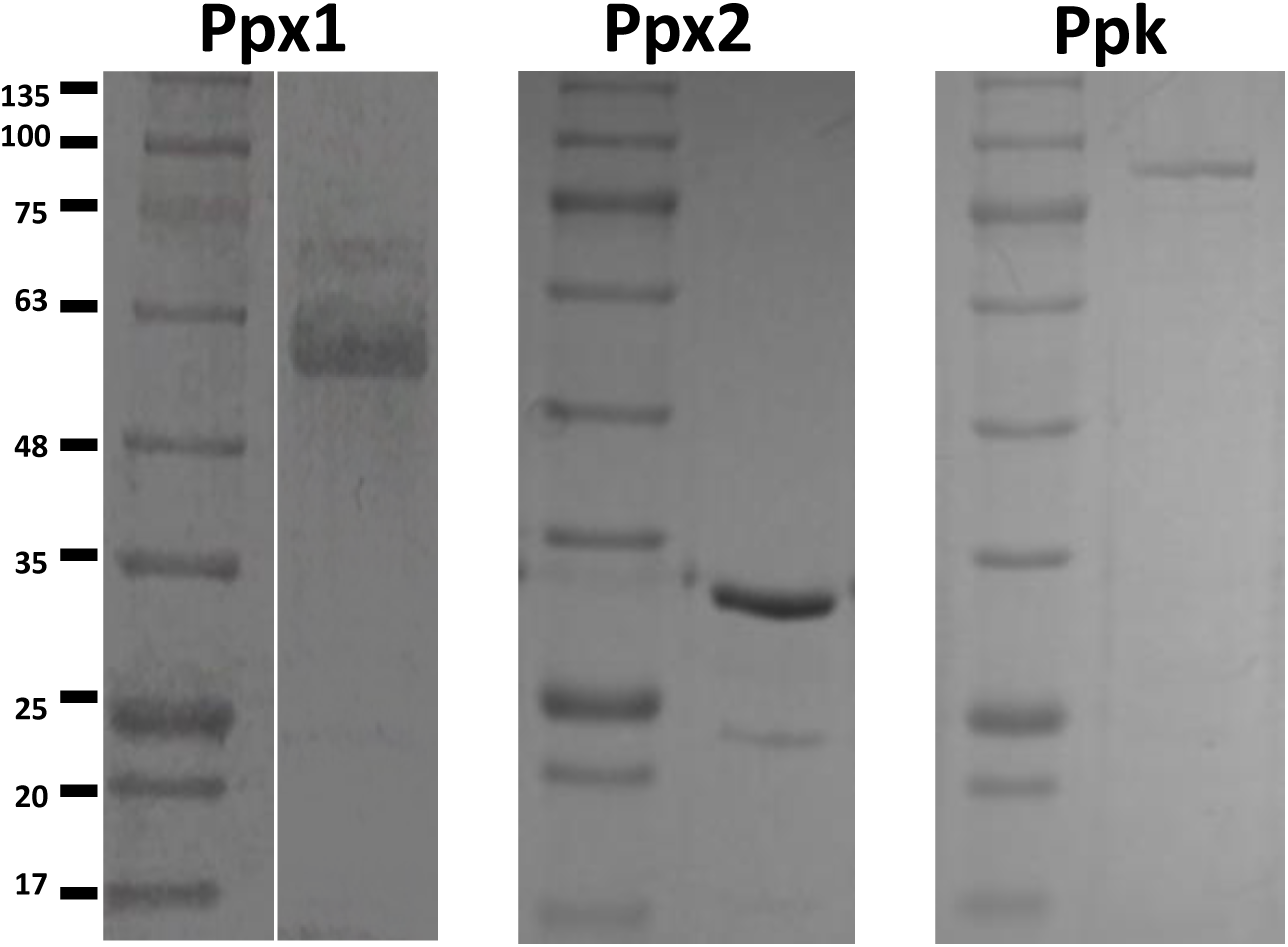
PAGE gels showing purified Ppx1 (57.3 kDa), Ppx2 (35.3 kDa) and Ppk (82.7 KDa) from *Lc. paracasei* BL23. The molecular weight reference is the NZYColour Molecular Weight Marker II.

**Supplementary Figure S2.**
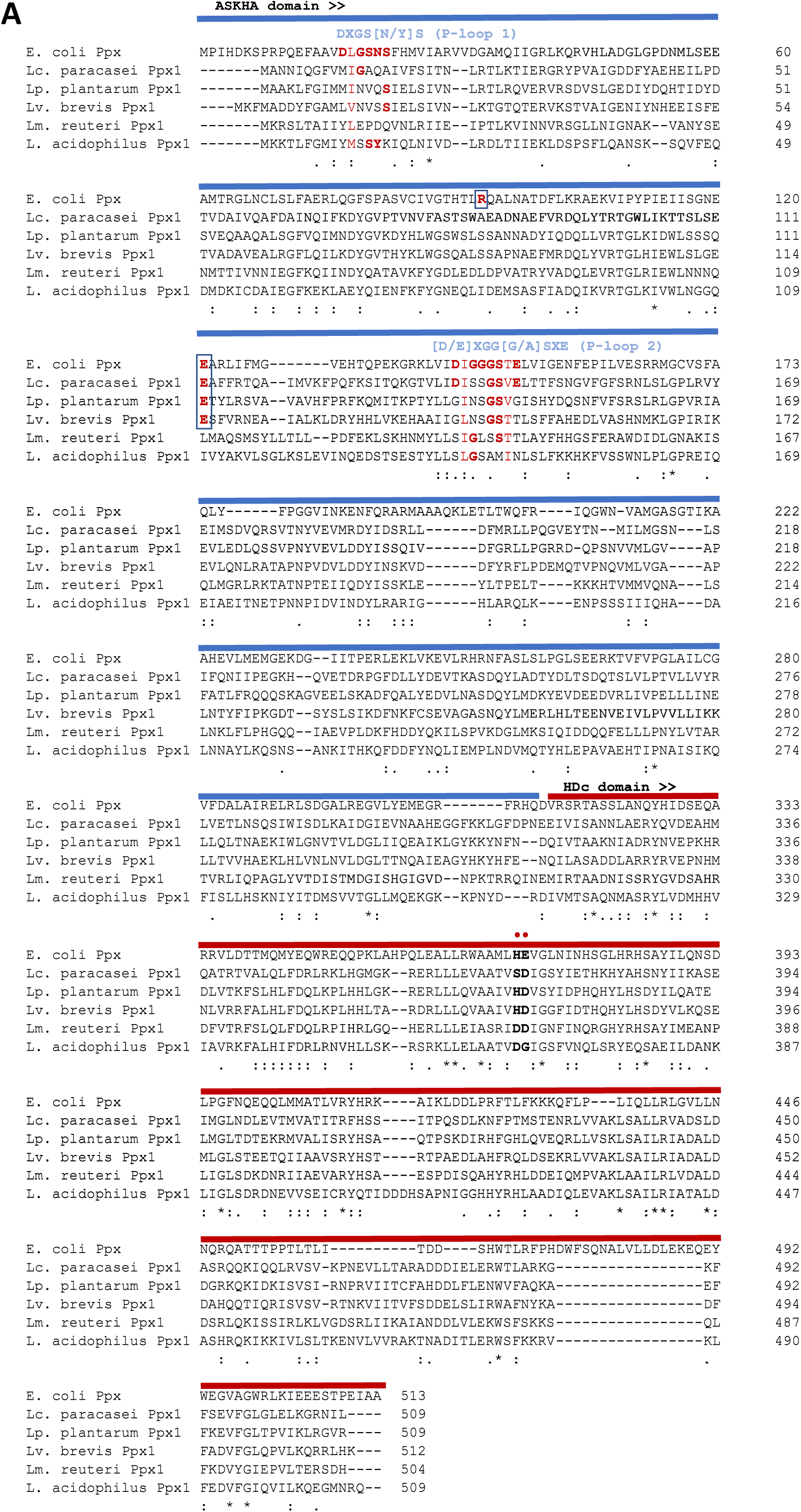
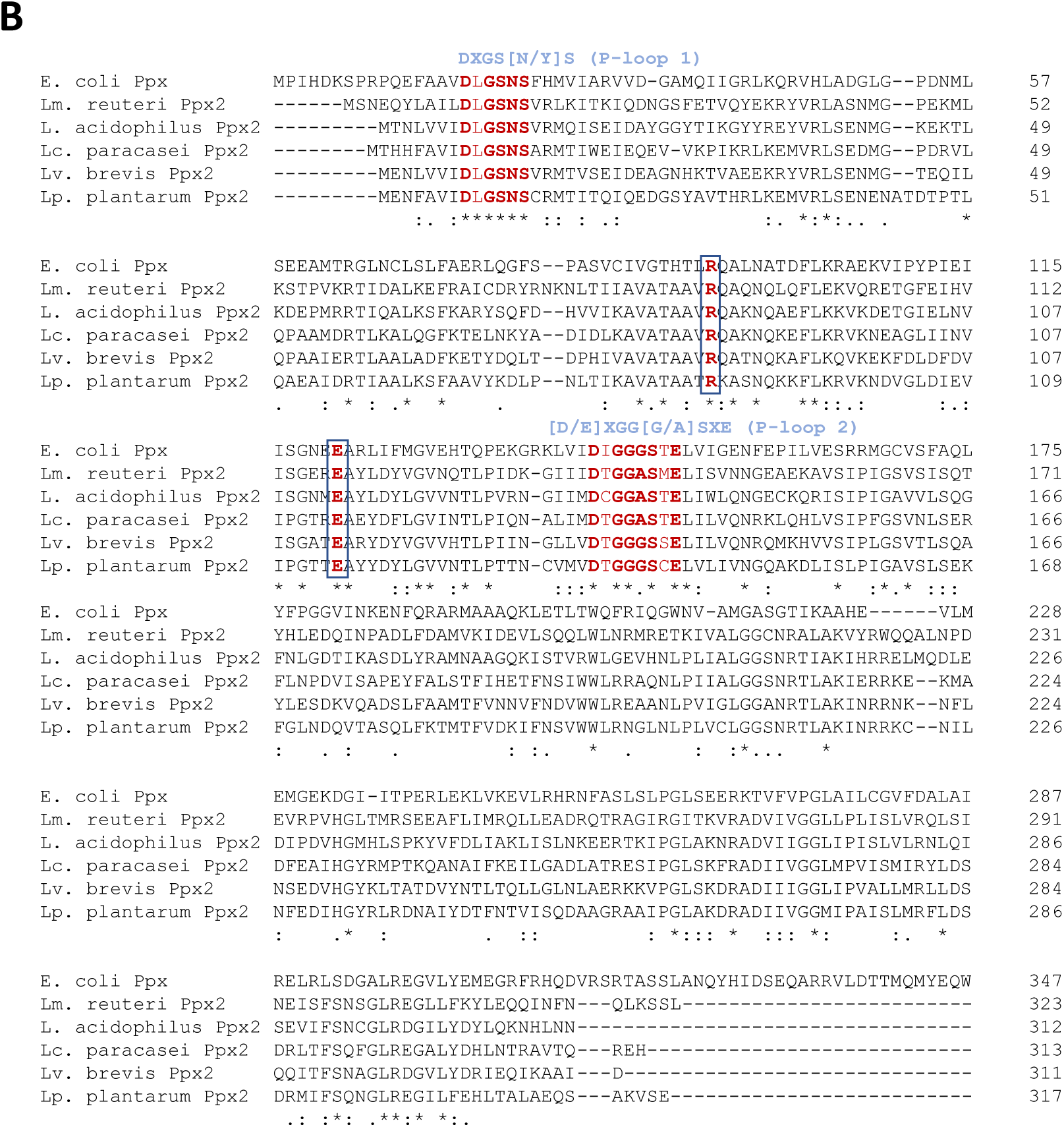
Multiple amino acid alignments of Ppx proteins from members of the Lactobacillaceae. (A) Ppx1 from *Lc. paracasei* (WP_003606084.1), *Lv. brevis* (AYM03703.1), *Lp. plantarum* (WP_103851489.1), *Lm. reuteri* (MCH5356744.1) and *L. acidophilus* (MCT3601606.1) (B) Ppx2 from *Lc. paracasei* (CAQ67832.1), *Lv. brevis* (WP_135367400.1), *Lp. plantarum* (WP_003641094.1), *Lm. reuteri* (KRK50994.1) and *L. acidophilus* (AZN76677.1). Ppx from *E. coli* (WP_001121363.1) was included in both alignments for comparison. In the Ppx2 alignment, only the first 347 amino acids from *E. coli* Ppx are shown. The consensus sequences of the conserved P-loop 1 and P-loop 2 in the GppA/Ppx family are shown in blue above the alignments. Amino acids in bold show those matching that consensus. The boxed amino acids indicate the glutamic and arginine residues involved in catalysis in the GppA/Ppx family. The ASKHA (acetate and sugar kinase/Hsp70/actin) and HDc domains in Ppx1 are indicated in blue and red, respectively. The amino acids characteristic of the HDc domain in C-terminal Ppx1 are marked with red circles.

## REFERENCES

1. Albi T, Serrano A. 2014. Two exopolyphosphatases with distinct molecular architectures and substrate specificities from the thermophilic green-sulfur bacterium Chlorobium tepidum TLS. Microbiology (Reading) 160:2067–2078.

2. Amado L, Kuzminov A. 2009. Polyphosphate accumulation in Escherichia coli in response to defects in DNA metabolism. J Bacteriol 191:7410–7416.

3. Gray MJ, Jakob U. 2015. Oxidative stress protection by polyphosphate--new roles for an old player. Curr Opin Microbiol 24:1–6.

4. Candon HL, Allan BJ, Fraley CD, Gaynor EC. 2007. Polyphosphate kinase 1 is a pathogenesis determinant in Campylobacter jejuni. J Bacteriol 189:8099–8108.

5. Malde A, Gangaiah D, Chandrashekhar K, Pina-Mimbela R, Torrelles JB, Rajashekara G. 2014. Functional characterization of exopolyphosphatase/guanosine pentaphosphate phosphohydrolase (PPX/GPPA) of Campylobacter jejuni. Virulence 5:521–533.

6. Peng L, Jiang Q, Pan JY, Deng C, Yu JY, Wu XM, Huang SH, Deng XY. 2016. Involvement of polyphosphate kinase in virulence and stress tolerance of uropathogenic Proteus mirabilis. Med Microbiol Immunol 205:97–109.

7. Rashid MH, Rumbaugh K, Passador L, Davies DG, Hamood AN, Iglewski BH, Kornberg A. 2000. Polyphosphate kinase is essential for biofilm development, quorum sensing, and virulence of Pseudomonas aeruginosa. Proc Natl Acad Sci U S A 97:9636–9641.

8. Srisanga K, Suthapot P, Permsirivisarn P, Govitrapong P, Tungpradabkul S, Wongtrakoongate P. 2019. Polyphosphate kinase 1 of Burkholderia pseudomallei controls quorum sensing, RpoS and host cell invasion. J Proteomics 194:14–24.

9. Segawa S, Fujiya M, Konishi H, Ueno N, Kobayashi N, Shigyo T, Kohgo Y. 2011. Probiotic-derived polyphosphate enhances the epithelial barrier function and maintains intestinal homeostasis through integrin-p38 MAPK pathway. PLoS One 6:e23278.

10. Tanaka K, Fujiya M, Konishi H, Ueno N, Kashima S, Sasajima J, Moriichi K, Ikuta K, Tanabe H, Kohgo Y. 2015. Probiotic-derived polyphosphate improves the intestinal barrier function through the caveolin-dependent endocytic pathway. Biochem Biophys Res Commun 467:541–548.

11. Fujiya M, Ueno N, Kashima S, Tanaka K, Sakatani A, Ando K, Moriichi K, Konishi H, Kamiyama N, Tasaki Y, Omura T, Matsubara K, Taruishi M, Okumura T. 2020. Long-Chain Polyphosphate Is a Potential Agent for Inducing Mucosal Healing of the Colon in Ulcerative Colitis. Clin Pharmacol Ther 107:452–461.

12. Isozaki S, Konishi H, Fujiya M, Tanaka H, Murakami Y, Kashima S, Ando K, Ueno N, Moriichi K, Okumura T. 2021. Probiotic-Derived Polyphosphate Accelerates Intestinal Epithelia Wound Healing through Inducing Platelet-Derived Mediators. Mediators Inflamm 2021:5582943.

13. Saiki A, Ishida Y, Segawa S, Hirota R, Nakamura T, Kuroda A. 2016. A Lactobacillus mutant capable of accumulating long-chain polyphosphates that enhance intestinal barrier function. Biosci Biotechnol Biochem 80:955–961.

14. Rao NN, Gomez-Garcia MR, Kornberg A. 2009. Inorganic polyphosphate: essential for growth and survival. Annu Rev Biochem 78:605–647.

15. Ault-Riche D, Fraley CD, Tzeng CM, Kornberg A. 1998. Novel assay reveals multiple pathways regulating stress-induced accumulations of inorganic polyphosphate in Escherichia coli. J Bacteriol 180:1841–1847.

16. Kuroda A, Murphy H, Cashel M, Kornberg A. 1997. Guanosine tetra- and pentaphosphate promote accumulation of inorganic polyphosphate in Escherichia coli. J Biol Chem 272:21240–21243.

17. Racki LR, Tocheva EI, Dieterle MG, Sullivan MC, Jensen GJ, Newman DK. 2017. Polyphosphate granule biogenesis is temporally and functionally tied to cell cycle exit during starvation in Pseudomonas aeruginosa. Proc Natl Acad Sci U S A 114:E2440–E2449.

18. Gray MJ. 2019. Inorganic Polyphosphate Accumulation in Escherichia coli Is Regulated by DksA but Not by (p)ppGpp. J Bacteriol 201.

19. Bowlin MQ, Long AR, Huffines JT, Gray MJ. 2022. The role of nitrogen-responsive regulators in controlling inorganic polyphosphate synthesis in Escherichia coli. Microbiology (Reading) 168.

20. Gray MJ. 2020. Interactions between DksA and Stress-Responsive Alternative Sigma Factors Control Inorganic Polyphosphate Accumulation in Escherichia coli. J Bacteriol 202.

21. Alcantara C, Blasco A, Zuniga M, Monedero V. 2014. Accumulation of polyphosphate in Lactobacillus spp. and its involvement in stress resistance. Appl Environ Microbiol 80:1650–1659.

22. Alcantara C, Coll-Marques JM, Jadan-Piedra C, Velez D, Devesa V, Zuniga M, Monedero V. 2018. Polyphosphate in Lactobacillus and Its Link to Stress Tolerance and Probiotic Properties. Front Microbiol 9:1944.

23. Pokhrel A, Lingo JC, Wolschendorf F, Gray MJ. 2019. Assaying for Inorganic Polyphosphate in Bacteria. J Vis Exp doi:10.3791/58818.

24. Leloup L, Ehrlich SD, Zagorec M, Morel-Deville F. 1997. Single-crossover integration in the Lactobacillus sake chromosome and insertional inactivation of the ptsI and lacL genes. Appl Environ Microbiol 63:2117–2123.

25. Posno M, Leer RJ, van Luijk N, van Giezen MJ, Heuvelmans PT, Lokman BC, Pouwels PH. 1991. Incompatibility of Lactobacillus Vectors with Replicons Derived from Small Cryptic Lactobacillus Plasmids and Segregational Instability of the Introduced Vectors. Appl Environ Microbiol 57:1822–1828.

26. Schotte L, Steidler L, Vandekerckhove J, Remaut E. 2000. Secretion of biologically active murine interleukin-10 by Lactococcus lactis. Enzyme Microb Technol 27:761–765.

27. Aschar-Sobbi R, Abramov AY, Diao C, Kargacin ME, Kargacin GJ, French RJ, Pavlov E. 2008. High sensitivity, quantitative measurements of polyphosphate using a new DAPI-based approach. J Fluoresc 18:859–866.

28. Ghorbel S, Smirnov A, Chouayekh H, Sperandio B, Esnault C, Kormanec J, Virolle MJ. 2006. Regulation of ppk expression and in vivo function of Ppk in Streptomyces lividans TK24. J Bacteriol 188:6269–6276.

29. Zuniga M, Miralles Md Mdel C, Perez-Martinez G. 2002. The Product of arcR, the sixth gene of the arc operon of Lactobacillus sakei, is essential for expression of the arginine deiminase pathway. Appl Environ Microbiol 68:6051–6058.

30. Landete JM, Garcia-Haro L, Blasco A, Manzanares P, Berbegal C, Monedero V, Zuniga M. 2010. Requirement of the Lactobacillus casei MaeKR two-component system for L-malic acid utilization via a malic enzyme pathway. Appl Environ Microbiol 76:84–95.

31. Pfaffl MW, Horgan GW, Dempfle L. 2002. Relative expression software tool (REST) for group-wise comparison and statistical analysis of relative expression results in real-time PCR. Nucleic Acids Res 30:e36.

32. Jagannathan V, Kaur P, Datta S. 2010. Polyphosphate kinase from M. tuberculosis: an interconnect between the genetic and biochemical role. PLoS One 5:e14336.

33. Song H, Dharmasena MN, Wang C, Shaw GX, Cherry S, Tropea JE, Jin DJ, Ji X. 2020. Structure and activity of PPX/GppA homologs from Escherichia coli and Helicobacter pylori. FEBS J 287:1865–1885.

34. Alvarado J, Ghosh A, Janovitz T, Jauregui A, Hasson MS, Sanders DA. 2006. Origin of exopolyphosphatase processivity: Fusion of an ASKHA phosphotransferase and a cyclic nucleotide phosphodiesterase homolog. Structure 14:1263–1272.

35. Zhang A, Lu Z, Xu Y, Qi T, Li W, Zhang L, Cui Z. 2021. The structure of exopolyphosphatase (PPX) from Porphyromonas gingivalis in complex with substrate analogs and magnesium ions reveals the basis for polyphosphate processivity. J Struct Biol 213:107767.

36. Correa Deza MA, Grillo-Puertas M, Salva S, Rapisarda VA, Gerez CL, Font de Valdez G. 2017. Inorganic salts and intracellular polyphosphate inclusions play a role in the thermotolerance of the immunobiotic Lactobacillus rhamnosus CRL 1505. PLoS One 12:e0179242.

37. Munevar NF, de Almeida LG, Spira B. 2017. Differential regulation of polyphosphate genes in Pseudomonas aeruginosa. Mol Genet Genomics 292:105–116.

38. de Almeida LG, Ortiz JH, Schneider RP, Spira B. 2015. phoU inactivation in Pseudomonas aeruginosa enhances accumulation of ppGpp and polyphosphate. Appl Environ Microbiol 81:3006–3015.

39. Morohoshi T, Maruo T, Shirai Y, Kato J, Ikeda T, Takiguchi N, Ohtake H, Kuroda A. 2002. Accumulation of inorganic polyphosphate in phoU mutants of Escherichia coli and Synechocystis sp. strain PCC6803. Appl Environ Microbiol 68:4107–4110.

40. Geissdorfer W, Ratajczak A, Hillen W. 1998. Transcription of ppk from Acinetobacter sp. strain ADP1, encoding a putative polyphosphate kinase, is induced by phosphate starvation. Appl Environ Microbiol 64:896–901.

41. Gallarato LA, Sanchez DG, Olvera L, Primo ED, Garrido MN, Beassoni PR, Morett E, Lisa AT. 2014. Exopolyphosphatase of Pseudomonas aeruginosa is essential for the production of virulence factors, and its expression is controlled by NtrC and PhoB acting at two interspaced promoters. Microbiology (Reading) 160:406–417.

42. Silby MW, Nicoll JS, Levy SB. 2009. Requirement of polyphosphate by Pseudomonas fluorescens Pf0-1 for competitive fitness and heat tolerance in laboratory media and sterile soil. Appl Environ Microbiol 75:3872–3881.

43. Rudat AK, Pokhrel A, Green TJ, Gray MJ. 2018. Mutations in Escherichia coli Polyphosphate Kinase That Lead to Dramatically Increased In Vivo Polyphosphate Levels. J Bacteriol 200.

44. Lindner SN, Knebel S, Wesseling H, Schoberth SM, Wendisch VF. 2009. Exopolyphosphatases PPX1 and PPX2 from Corynebacterium glutamicum. Appl Environ Microbiol 75:3161–3170.

45. Nikel PI, Chavarria M, Martinez-Garcia E, Taylor AC, de Lorenzo V. 2013. Accumulation of inorganic polyphosphate enables stress endurance and catalytic vigour in Pseudomonas putida KT2440. Microb Cell Fact 12:50.

46. Aravind L, Koonin EV. 1998. The HD domain defines a new superfamily of metal-dependent phosphohydrolases. Trends Biochem Sci 23:469–472.

47. Bolesch DG, Keasling JD. 2000. Polyphosphate binding and chain length recognition of Escherichia coli exopolyphosphatase. J Biol Chem 275:33814–33819.

48. Choi MY, Wang Y, Wong LL, Lu BT, Chen WY, Huang JD, Tanner JA, Watt RM. 2012. The two PPX-GppA homologues from Mycobacterium tuberculosis have distinct biochemical activities. PLoS One 7:e42561.

49. Beassoni PR, Gallarato LA, Boetsch C, Garrido MN, Lisa AT. 2015. Pseudomonas aeruginosa Exopolyphosphatase Is Also a Polyphosphate: ADP Phosphotransferase. Enzyme Res 2015:404607.

50. Chawla R, Klupt S, Patsalo V, Williamson JR, Racki LR. 2022. The Histone H1-Like Protein AlgP Facilitates Even Spacing of Polyphosphate Granules in Pseudomonas aeruginosa. mBio 13:e0246321.

51. Tumlirsch T, Jendrossek D. 2017. Proteins with CHADs (Conserved Histidine alpha-Helical Domains) Are Attached to Polyphosphate Granules In Vivo and Constitute a Novel Family of Polyphosphate-Associated Proteins (Phosins). Appl Environ Microbiol 83.

52. Tumlirsch T, Sznajder A, Jendrossek D. 2015. Formation of polyphosphate by polyphosphate kinases and its relationship to poly(3-hydroxybutyrate) accumulation in Ralstonia eutropha strain H16. Appl Environ Microbiol 81:8277–8293.

53. Henry JT, Crosson S. 2013. Chromosome replication and segregation govern the biogenesis and inheritance of inorganic polyphosphate granules. Mol Biol Cell 24:3177–3186.

54. Maze A, Boel G, Zuniga M, Bourand A, Loux V, Yebra MJ, Monedero V, Correia K, Jacques N, Beaufils S, Poncet S, Joyet P, Milohanic E, Casaregola S, Auffray Y, Perez-Martinez G, Gibrat JF, Zagorec M, Francke C, Hartke A, Deutscher J. 2010. Complete genome sequence of the probiotic Lactobacillus casei strain BL23. J Bacteriol 192:2647–2648.

